# The impact of cigarette smoke exposure, and COPD or asthma status on ABC transporter gene expression in human airway epithelial cells

**DOI:** 10.1101/369355

**Authors:** Jennifer A. Aguiar, Andrea Tamminga, Briallen Lobb, Ryan D. Huff, Jenny Nguyen, Yechan Kim, Anna Dvorkin-Gheva, Martin R. Stampfli, Andrew C. Doxey, Jeremy A. Hirota

**Affiliations:** Department of Biology, University of Waterloo, Waterloo, ON, Canada N2L 3G1; Division of Respiratory Medicine, Department of Medicine, University of British Columbia, Vancouver, BC, Canada, V6H 3Z6; Firestone Institute for Respiratory Health – Division of Respirology, Department of Medicine, McMaster University, Hamilton, ON, L8N 4A6; McMaster Immunology Research Centre, McMaster University, Hamilton, ON, Canada, L8S 4K1; Department of Medicine, McMaster University, Hamilton, ON, Canada, L8S 4K1

## Abstract

**Rationale:** The respiratory mucosa coordinates responses to infections, allergens, and exposures to air pollution. A relatively unexplored aspect of the respiratory mucosa are the expression and function of ATP Binding Cassette (ABC) transporters. ABC transporters are conserved in prokaryotes and eukaryotes, with humans expressing 48 transporters divided into 7 classes (ABCA, ABCB, ABCC, ABCD, ABDE, ABCF, and ABCG). Throughout the human body, ABC transporters regulate cAMP levels, chloride secretion, lipid transport, and anti-oxidant responses. A deeper exploration of the expression patterns of ABC transporters in the respiratory mucosa is warranted to determine their relevance in lung health and disease.

**Methods:** We used a bioinformatic approach complemented with *in vitro* experimental methods for validation of candidate ABC transporters. We analyzed the expression profiles of all 48 human ABC transporters in the respiratory mucosa using bronchial epithelial cell gene expression datasets available in NCBI GEO from well-characterized patient populations of healthy subjects and individuals that smoke cigarettes, or have been diagnosed with COPD or asthma. The Calu-3 airway epithelial cell line was used to interrogate selected results using a cigarette smoke extract exposure model.

**Results:** Using 9 distinct gene-expression datasets of primary human airway epithelial cells, we completed a focused analysis on 48 ABC transporters in samples from healthy subjects and individuals that smoke cigarettes, or have been diagnosed with COPD or asthma. *In situ* gene expression data demonstrate that ABC transporters are i) variably expressed in epithelial cells from different airway generations (top three expression levels - *ABCA5, ABCA13*, and *ABCC5)*, ii) regulated by cigarette smoke exposure (*ABCA13*, *ABCB6*, *ABCC1*, and *ABCC3*), and iii) differentially expressed in individuals with COPD and asthma (*ABCA13*, *ABCC1*, *ABCC2*, *ABCC9*). An *in vitro* cell culture model of cigarette smoke exposure was able to recapitulate the *in situ* changes observed in cigarette smokers for *ABCA13* and *ABCC1*.

**Conclusions:** Our *in situ* human gene expression data analysis reveals that ABC transporters are expressed throughout the airway generations in airway epithelial cells and can be modulated by environmental exposures important in chronic respiratory disease (e.g. cigarette smoking) and in individuals with chronic lung diseases (e.g. COPD or asthma). Our work highlights select ABC transporter candidates of interest and a relevant *in vitro* model that will enable a deeper understanding of the contribution of ABC transporters in the respiratory mucosa in lung health and disease.

## Introduction

The respiratory mucosa, extending from the upper airways down to simple squamous epithelium, provides a continuous physical barrier with immune function that helps protect us from any potentially harmful substances contained in the air we breathe^1,2^. The many functions of the respiratory mucosa include the ability to produce cytokines to coordinate immune responses, regulate ion concentrations for airway surface lining fluid and mucus function, and detect viral entry into epithelial cells for antiviral immune responses. In health, these coordinated functions can ensure that infections, allergens, and exposures to air pollution are controlled to limit damage to host, while in disease, a dysfunction in any capacity of the respiratory mucosa may lead to development or exacerbation of chronic respiratory disease. It is therefore important to understand the mechanisms that regulate respiratory mucosa function.

A relatively unexplored contributor to respiratory mucosa biology is the ATP Binding Cassette (ABC) family of transporters^3-7^. ABC transporters are conserved in prokaryotes and eukaryotes, with humans expressing 48 transporters divided into 7 classes (ABCA, ABCB, ABCC, ABCD, ABDE, ABCF, and ABCG)(7). The majority of ABC transporters couple ATP hydrolysis to the extracellular transport of substrates. ABC transporters have been demonstrated to transport cytokines, ions, lipids, as well as detect viral insults, although the confirmation of these functions and expression of these genes and protein has not been extensively confirmed in the respiratory mucosa of humans^8,9^. Our overarching hypothesis is that ABC transporter expression in human airway epithelial cells contributes to respiratory mucosal immunity that modulates the lung environment in response to environmental exposures including cigarette smoke, allergens, air pollution, bacteria, and viruses.

The precedent for the importance of ABC transporters in respiratory mucosa biology has been set by the identification of mutations in ABC transporter genes in patients with lung pathologies. ABCC7, also known as cystic fibrosis transmembrane conductance regulator (CFTR), is the causal gene contributing to the development of cystic fibrosis^10,11^. Over 2000 variants in CFTR have been reported, resulting in altered chloride and bicarbonate secretion, hydration of the airway surface lining fluid, and mucus function^10,11^. Equally convincing of the role of ABC transporters in lung health and disease is the link between genetic loss of function variants of ABCA3 and fatal surfactant deficiency in newborns^12^. In addition, our group has recently confirmed that ABCC4 is expressed in airway epithelial cells^4,5^, is able to transport cAMP and uric acid^5^, and can modulate anti-inflammatory activities of long-acting beta-agonist/glucocorticoid therapies^6^ and potentiate CFTR in select patient populations^13^. These three examples represent only a fraction of the 48 transporters that remain to be more extensively studied in the respiratory mucosa for expression patterns and function in lung health and disease.

We have initiated a characterization of all 48 ABC transporters in the respiratory mucosa by performing a bioinformatic analysis of 9 distinct gene-expression datasets of primary human airway epithelial cells isolated from bronchial brushings of well-phenotyped healthy subjects and individuals that smoke cigarettes, or have been diagnosed with COPD or asthma^14-22^. Our hypothesis was that specific ABC transporter gene expression patterns would correlate with presence of a specific chronic respiratory disease and disease severity. To validate our bioinformatic analyses, we interrogated select results using an *in vitro* cell culture model system to provide a foundational platform for further interrogation of candidate ABC transporters identified that may be important in chronic respiratory disease pathology. Our *in situ* human gene expression data analysis reveals that ABC transporters are i) variably expressed in epithelial cells from different airway generations, ii) regulated by cigarette smoke exposure, iii) differentially expressed in individuals with COPD and asthma, and iv) are amenable to interrogation with *in vitro* cell culture systems. We conclude that continued research into the basic biology of ABC transporters in the respiratory mucosa is required to truly define the importance of these molecules in lung health and disease.

## Methods

### Dataset selection and quality control

Analysis was performed on previously deposited datasets obtained from the Gene Expression Omnibus (GEO). Nine datasets were used during the course of this study. Datasets pertaining to smoking and COPD analysis include GSE994, GSE4498, GSE11784, GSE11906, and GSE37147. Datasets focusing on asthma and severity include GSE4302, GSE63142, GSE67472, and GSE76227^14-22^.

Since the datasets were generated from a variety of studies, microarray chip selection and data normalization methods vary across the datasets. GSEs 11906, 11784, 4498, and 994 use the MAS5 normalization method without log transformation, whereas GSEs 76227, 4302, 67472, and 37147 use the Robust Multi-array Average (RMA) method with log transformation. GSE63142 uses yet another normalization method, cyclic LOESS. To avoid applying multiple compounding normalization methods to any given dataset or attempting to reverse already applied normalizations, datasets were left as provided by the original studies and differences in methods were noted throughout our analysis and in figure legends. Since probeset IDs and corresponding gene targets vary across microarray platforms, expression results are reported in the context of both ABC transporter name and probeset ID.

### ABC transporter transcript expression in human airway epithelial cells

Differential expression patterns for all 48 ABC transporter members were examined across the trachea, large airways (generation 2^nd^-3^rd^), and small airways (generation 10^th^-12^th^) using a single dataset (GSE11906, Affymetrix Human Genome U133 Plus 2 microarray platform)^17^. A bar plot showing average expression of ABC transporters in healthy, non-smoker lung tissue from GSE11906 was generated using the ggplot2 package in R (v. 3.4.3) with values categorized by airway location. In GSE11906, independent samples of epithelial cells were isolated from healthy individuals with no smoking history and normal lung function and included 17 trachea (age – 42 +/− 7), 21 large airway (age – 42 +/− 9), and 35 small airway samples(15) (age 43 +/− 10).

### Impact of cigarette smoke exposure on ABC transporter expression

Three datasets (GSE11906, GSE11784, and GSE4498), which were all generated from the Affymetrix Human Genome U133 Plus 2 microarray platform, were used to assess the impact of cigarette smoke exposure on ABC transporter gene expression profiles^15-17^. All three datasets are comprised of small airway (10^th^-12^th^ generation) epithelial cell transcript expression patterns in healthy subjects and those with a history of smoking without a diagnosis of COPD. In GSE11906, 54 independent samples of epithelial cells were isolated from individuals with >25 pack years smoking history with no reported COPD^17^. In GSE11784, 72 independent samples of epithelial cells were isolated from individuals with >25 pack years smoking history with no reported COPD^16^. In GSE4498, 10 independent samples of epithelial cells were isolated from individuals with >25 pack years smoking history with no reported COPD^15^. We independently curated these three GSE datasets to ensure that no samples were repeated across the analyses.

### Impact of cigarette smoking cessation on ABC transporter expression

Two datasets (GSE37147 and GSE994) generated from two distinct microarray platforms (Affymetrix Human Gene 1 ST and Affymetrix Human Genome U133A, respectively), which analyzed epithelial cells from medium (6^th^-8^th^ generation) and large (2^nd^ generation) airways, respectively, were used to assess the impact of smoking cessation on ABC transporter gene expression profiles^14,18^. In GSE37147, independent samples of epithelial cells were isolated from 69 current smokers and 82 former smokers with >47 pack years smoking history with no reported COPD^18^. In GSE37147, the average duration of smoking cessation was 11.11 years. In GSE994, independent samples of epithelial cells were isolated from 34 current smokers, 14 former smokers, and 23 never-smokers, with >22 pack years smoking history with no reported COPD^14^. In GSE994, the average duration of smoking cessation was 10.49 years.

### Association between COPD status and changes in ABC transporter expression

Three datasets (GSE11906, GSE11784, and GSE37147) were used to assess association of COPD status with ABC transporter gene expression profile. GSE11906 and GSE11784 collected epithelial cells from small airways (10^th^-12^th^ generation) while GSE37147 collected from medium airways (6^th^-8^th^ generation), with the two different sample types also analyzed on different microarray platforms (Affymetrix Human Genome U133 Plus 2 and Affymetrix Human Gene 1 ST, respectively).

In GSE11906, 20 independent samples of epithelial cells were isolated from individuals with >38 pack years smoking history with reported COPD and were compared to 54 independent samples of epithelial cells isolated from individuals with >25 pack years smoking history with no reported COPD^17^. In GSE11784, 36 independent samples of epithelial cells were isolated from individuals with >34 pack years smoking history with reported COPD and were compared to 72 independent samples of epithelial cells isolated from individuals with >25 pack years smoking history with no reported COPD^16^. In GSE37147, 87 independent samples of epithelial cells were isolated from individuals with >51 pack years smoking history with reported COPD and were compared to 151 independent samples of epithelial cells isolated from individuals with >47 pack years smoking history with no reported COPD^18^.

### ABC transporter expression patterns in airway epithelial cells from asthmatics

We analyzed ABC transporter expression in airway epithelial cells isolated from asthmatics relative to healthy controls. For our asthma-focused analysis we interrogated two pairs of datasets: GSE4302 and GSE67472 ^19-21^ – which allowed for healthy control comparison to individuals with asthma and GSE63142 and GSE76227 ^20-22^ – which allowed for associating expression profiles with asthma severity. All studies analyzed epithelial cells from medium (3^rd^-5^th^ generation) airways, while distinct microarray platforms were used (GSE4302 and GSE67472 used Affymetrix Human Genome U133 Plus 2, GSE63142 used Agilent 014850 Whole Human Genome Microarray 4×44K G4112F, and GSE76227 used Affymetrix HT HG U133 plus PM).

In GSE4302, independent samples of epithelial cells were isolated from 28 healthy individuals and 42 asthmatics^19^. In GSE67472, independent samples of epithelial cells were isolated from 43 healthy individuals and 62 asthmatics^21^. In GSE63142, independent samples of epithelial cells were isolated from 26 healthy individuals, 59 mild asthmatics, 19 moderate asthmatics and 51 severe asthmatics^20^. In GSE76227, independent samples of epithelial cells were isolated from 26 healthy individuals, 59 mild asthmatics, 19 moderate asthmatics and 51 severe asthmatics^22^.

### *In vitro* validation of candidate ABC transporter gene expression changes

A cigarette smoke extract conditioned media experiment with the Calu-3 airway epithelial cell line grown under submerged monolayer conditions was used for *in vitro* validation of candidate ABC transporters identified in our analysis of epithelial brushings. We performed an exposure with cigarette smoke extract conditioned media (10% and 20%) for 24hrs as previously described ^23^ and assessed *ABCA13* (forward: 5’-TGTGCTATTGTAACTCCTCTGAGAC-3’, reverse 5’-AAGTAATGACCTCTTGCAAAATACG-3’) and *ABCC1* (forward: 5’-GCCTATTACCCCAGCATCG-3’, reverse 5’-GATGCAGTTGCCCACACA-3’) gene expression relative to *GAPDH* (forward: 5’-ACGGGAAGCTTGTCATCAAT-3’ Reverse: 5’-CATCGCCCCACTTGATTTT-3’) by quantitative-PCR.

### Statistical analyses

Datasets were pruned to only contain the data matrix (probeset ID as column 1 and sample IDs populating subsequent columns) and relevant categorical data such as smoking status, disease status or airway location from which samples were isolated. Complete datasets including all qualitative information were saved separately for reference.

Lists of probeset IDs for each microarray platform used in the study were generated using only probes that corresponded to ABC transporter proteins. These lists were then used to filter corresponding matrices to include data exclusively for probes that target ABC transporter proteins across samples in the datasets.

Pairwise Mann-Whitney U tests were performed to assess the statistical significance of differential gene expression between conditions. Adjusted *p*-values (*q*-values) were derived using the Benjamini Hochberg multiple hypothesis testing correction as implemented in R (v 3.4.3). This statistical method was chosen because *a priori* analysis of all dataset distributions showed that the data was non-Normal. Non-parametric methods such as Mann-Whitney U (which is the non-parametric equivalent of the unpaired Student’s T-test) make no assumptions about distribution and are therefore preferred for the analysis of non-Normal data.

Boxplots were generated for individual probesets (identified by probeset ID and corresponding ABC transporter protein) using the ggplot2 package in R (v. 3.4.3). Boxplots visualize the minimum value (bottom of vertical line), first to third quartile (box), median (horizontal line within box), and maximum value (top of vertical line) of a data distribution. To visualize the underlying raw data distributions, individual data values were plotted as points on top of each boxplot with randomized horizontal skew. For GSE994 which includes more than 2 independent categorical variables (current, former and never smokers) we used the Kruskal-Wallis test. The Kruskal-Wallis test is a non-parametric alternative to the one-way ANOVA test that does not assume a data distribution and is more robust to type II error than a simple pairwise test. The Tukey Honest Significant Difference (Tukey HSD) test was used to perform *post-hoc* pairwise comparisons with multiple testing corrections on probeset IDs found to be significantly differentially expressed across categories as determined by the Kruskal-Wallis analysis. To summarize the performed statistical analyses, tables were generated for each dataset that included Probeset ID, Probeset, the Benjamini Hochberg corrected *p*-value, and the log2 fold change. The top 10 probeset IDs were listed for each dataset based on corrected *p*-value, and multiple datasets used to analyze the same biological question were combined into final summary tables.

For the *in vitro* study of cigarette smoke extract exposure, an a one-way ANOVA was performed with a Bonferroni correction for multiple comparisons with a significance determined at p<0.05.

## Results

### ABC transporter transcript expression in human airway epithelial cells

To begin our characterization of the 48 known ABC transporters in airway epithelial cells, we examined expression patterns of each member across the trachea, large airways (generation 2^nd^-3^rd^), and small airways (generation 10^th^-12^th^) **(Figure 1 – Supplement Table 1)** using a single dataset (GSE11906). A global assessment of all ABC transporters across airway generations revealed diversity in their expression, with the highest detected expression levels for *ABCA5*, *ABCA13*, and *ABCC5* **(Figure 1)** and a trend for increased expression in the trachea relative to the small airways. Between the airway generations, differences were only observed between the small airways and the trachea (*q* < 0.05 – Supplement Table 1). The top three candidates ranked by fold-change in small airway relative to trachea were *ABCC1*, *ABCC4*, and *ABCB3* (all with *q* <0.05). Collectively the data suggest that ABC transporters are expressed in human airway epithelial cells with airway generation specific expression patterns.

**Figure 1.**
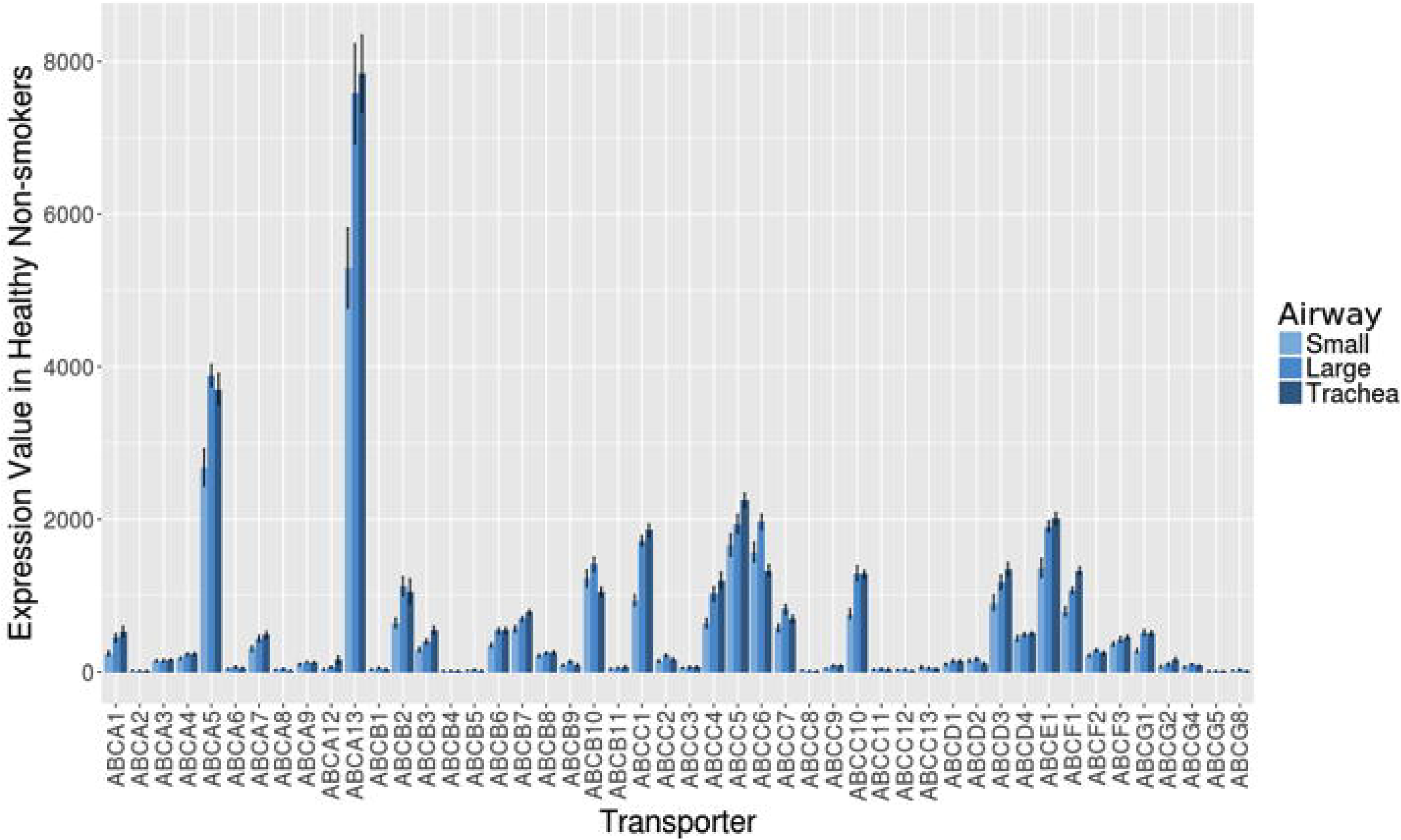
ABC transporter transcripts are expressed in airway epithelial cells throughout airway tree generations. Gene expression levels for all 48 ABC transporter members were examined across the trachea, large airways (generation 2^nd^-3^rd^), and small airways (generation 10^th^-12^th^) using a single dataset (GSE11906, Affymetrix Human Genome U133 Plus 2 microarray platform) comprised of independent epithelial cell samples isolated from healthy, non-smoking individuals. A representative probeset for each ABC transporter was chosen and average signal intensities (with standard error) for each probeset were calculated and visualized as a bar plot using ggplot2 in R (v. 3.4.3). Light blue = small airways, medium blue = large airways, dark blue = trachea.

### Impact of cigarette smoke exposure on ABC transporter expression

We next explored the impact of cigarette smoke exposure on ABC transporter expression profiles, as a model of environmental exposure with clinical relevance. For our analysis we interrogated three datasets from the same microarray platform of small airway (10^th^-12^th^ generation) epithelial cell transcript expression in healthy subjects and those with a history of smoking without a diagnosis of COPD (GSE11906, GSE11784, and GSE4498)^15-17^.

We observed that cigarette smoke exposure induced an increase in *ABCB6* and *ABCC3* expression levels, a feature conserved across all three independent datasets **(Figure 2 – Supplement Table 2).** In 2 of 3 datasets, an increase in *ABCC1* was observed, with similar non-significant trends observed in the third dataset. In 1 of 3 datasets a decrease in *ABCA13* was observed, with similar non-significant trends observed in the two other datasets.

**Figure 2.**
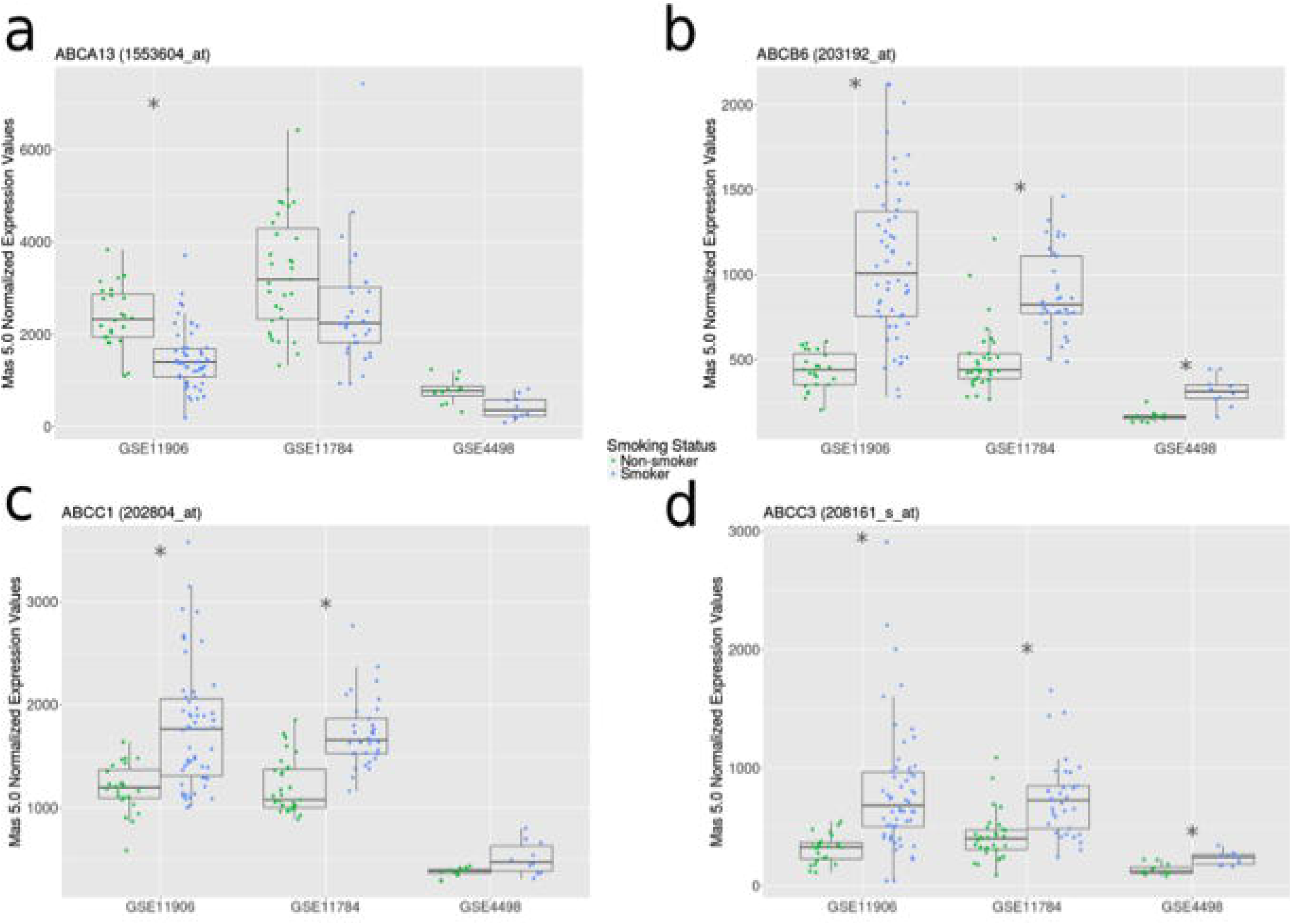
Cigarette smoking is associated with differential gene expression patterns of select ABC transporters in small airway epithelial cells. Gene expression levels of selected ABC transporters (*A*, *ABCA13*; *B*, *ABCB6*; *C*, *ABCC1*; *D*, *ABCC3*) between healthy non-smokers (green) and smokers with no diagnosis of COPD (blue). Significant expression differences are indicated by asterisks (*q* < 0.05 according to Mann-Whitney U test with Benjamini Hochberg correction). All three datasets used for this analysis (GSE11906, GSE11784, and GSE4498) analyzed epithelial cells from small airways (10^th^-12^th^ generation) and were generated from the Affymetrix Human Genome U133 Plus 2 microarray platform (using the Mas 5.0 Normalization method). Data distributions are visualized as boxplots (see Methods for additional details).

### Impact of cigarette smoking cessation on ABC transporter expression

Cigarette smoking is a modifiable risk factor for the development of COPD with cessation advocated by the Global Initiative for Obstructive Lung Disease^24^. We therefore sought to determine whether smoking cessation was able to normalize changes observed in ABC transporter expression for *ABCA13*, *ABCB6*, *ABCC1*, and *ABCC3* by comparing former smokers, current smokers, and where possible, healthy never-smokers. For our analysis we examined two datasets generated from two distinct microarray platforms, GSE37147 and GSE994, which analyzed epithelial cells from medium (6^th^-8^th^ generation) and large (2^nd^ generation) airways, respectively.

We observed that smoking cessation is associated with lower expression of *ABCB6*, *ABCC1*, and *ABCC3* in both datasets (**Figure 3 – Supplement Table 3**). A trend for elevated *ABCA13* expression (*q* = 0.06) was observed in association with smoking cessation in the medium airways with no primer sets available in the large airway dataset. In GSE994, an additional group of never-smokers reveals that levels of *ABCB6* and *ABCC1* elevated with smoking do not normalize to never-smoker levels even with prolonged smoking cessation (average 10.7 years)^14^.

**Figure 3.**
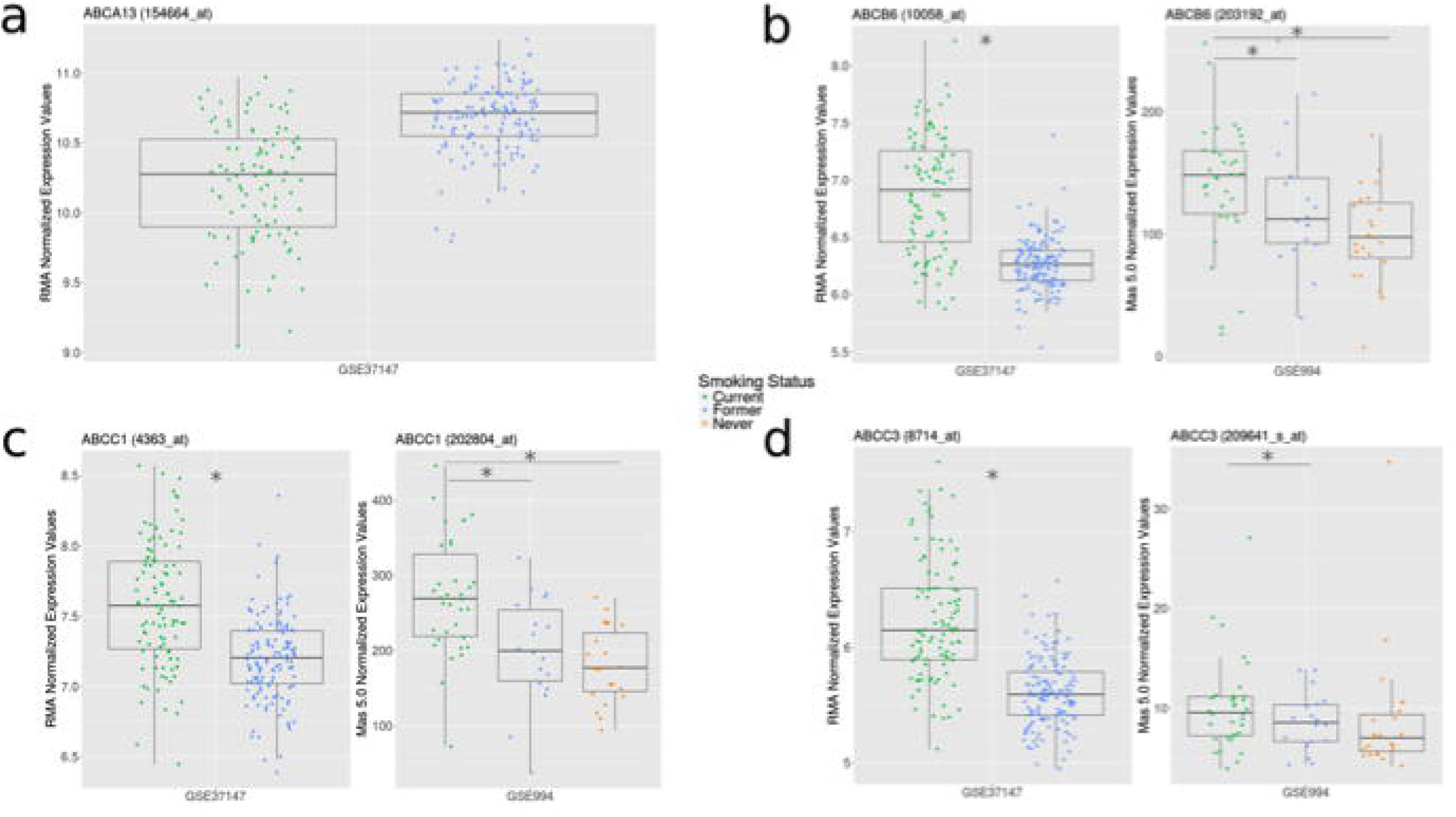
Cigarette smoking cessation is associated with normalization of ABC transporter gene expression patterns in airway epithelial cells. (A) Differential expression of *ABCA13* between current smokers (green) and former smokers (blue) from GSE37147 (* *q* < 0.05, Mann-Whitney U pairwise statistical analysis with Benjamini Hochberg multiple testing correction). GSE37147 was selected as it was the only dataset including a probe matching *ABCA13*. (B-D) Differential expression of *ABCB6*, *ABCC1*, *and ABCC3* between current smokers (green), former smokers (blue), and, where possible, never-smokers (orange) from GSE37147 and GSE994 (* *q* < 0.05, Kruskal-Wallis followed by the Tukey Honest Significance Difference (Tukey HSD) test). The datasets used in this analysis (GSE37147 and GSE994) were generated from two distinct microarray platforms (Affymetrix Human Gene 1 ST and Affymetrix Human Genome U133A, respectively), which analyzed epithelial cells from medium (6^th^-8^th^ generation) and large (2^nd^ generation) airways, respectively. GSE37147 used the logged RMA Normalization method and GSE994 used the Mas 5.0 Normalization method. Data distributions are visualized as boxplots (see Methods for additional details)

### Association between COPD status and changes in ABC transporter expression

We next analyzed the expression profiles of ABC transporters in relation to confirmed COPD status in three datasets (GSE11906, GSE11784, and GSE37147). GSE11906 and GSE11784 collected epithelial cells from small airways (10^th^-12^th^ generation) while GSE37147 collected from medium airways (6^th^-8^th^ generation), with the two different sample types also analyzed on different microarray platforms.

No ABC transporter showed differential expression across all three independent datasets (**Figure 4 – Supplement Table 4**). Of the candidates induced by cigarette smoke, *ABCC1* was ranked in the top 10 differentially expressed probesets in each dataset containing smokers with and without COPD, with a trend observed for increased expression with COPD status that was significant in the medium airways (GSE37147) (**Figure 4**). *ABCA13*, *ABCB6*, and *ABCC3* did not show any association with COPD status.

**Figure 4.**
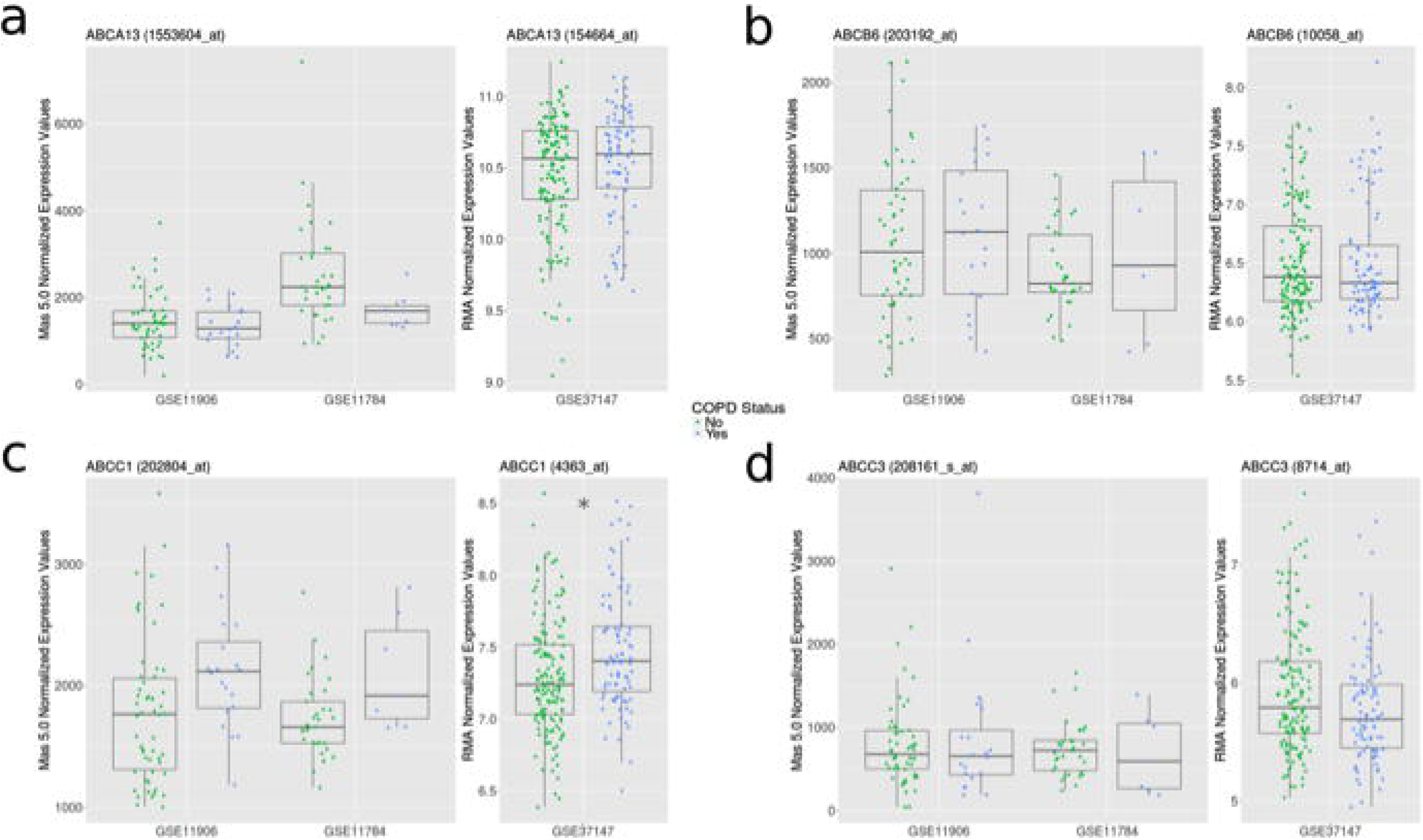
ABC transporter expression patterns associated with smoking status mildly correlate with COPD diagnosis. Gene expression levels of selected ABC transporters (*A*, *ABCA13*; *B*, *ABCB6*; *C*, *ABCC1*; *D*, *ABCC3*) between smokers with COPD (blue) and smokers with no diagnosis of COPD (green). Significant ABC transporter expression differences are indicated by asterisks (*q* < 0.05 according to Mann-Whitney U test with Benjamini Hochberg correction). Three datasets (GSE11906, GSE11784, and GSE37147) were used for this assessment. GSE11906 and GSE11784 collected epithelial cells from small airways (10^th^-12^th^ generation) while GSE37147 collected from medium airways (6^th^-8^th^ generation), with the two different sample types also analyzed on different microarray platforms (Affymetrix Human Genome U133 Plus 2 and Affymetrix Human Gene 1 ST, respectively). GSEs 11906 and 11784 used the Mas 5.0 Normalization method, and GSE37147 used the logged RMA Normalization method. Data distributions are visualized as boxplots (see Methods for additional details)

### ABC transporter expression patterns in airway epithelial cells from asthmatics

For our asthma-focused analysis we interrogated two pairs of datasets; GSE4302 and GSE67472 – which allowed for healthy control comparison to individuals with asthma and GSE63142 and GSE76227 – which allowed for associating expression profiles with asthma severity ^19-22^. All studies analyzed epithelial cells from medium (2^nd^-5^th^ generation) airways, while distinct microarray platforms were used.

We observed that asthma status is associated with an increase in *ABCC2* in both datasets (**Figure 5 – Supplement Table 5**). *ABCA13* and *ABCC9* were decreased in one dataset (GSE67472) with a similar trend in the second dataset (*q* < 0.1 – GSE4302). *ABCC1* expression was increased in one dataset (GSE4302) with no trend observed in the second dataset. *ABCC4* expression levels showed trends towards reduced expression in samples from asthmatics in both datasets that were not significant.

**Figure 5.**
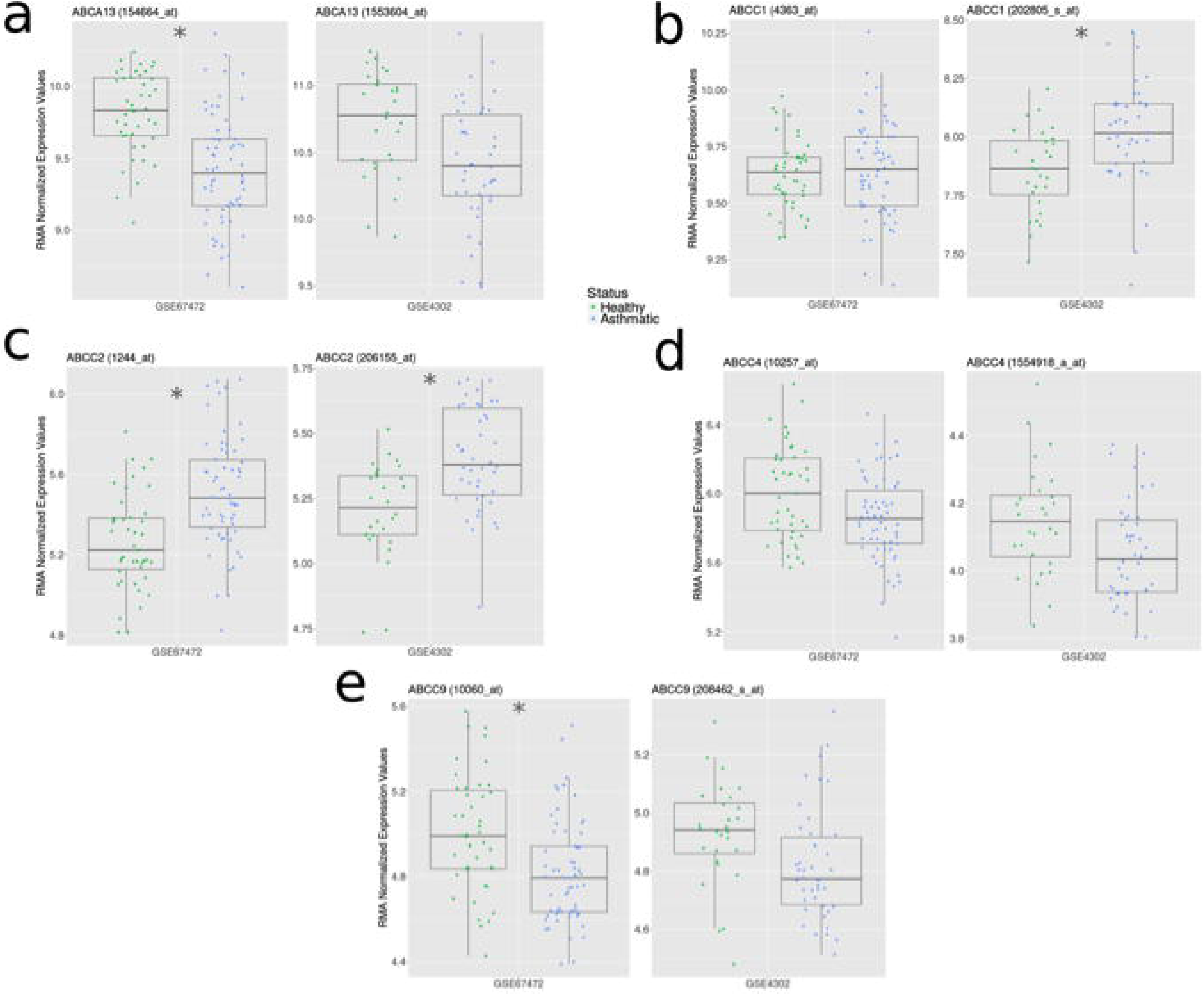
A change in ABC transporter expression patterns in airway epithelial cells from asthmatics is distinct from smoking or COPD status. Gene expression levels of selected ABC transporters (*A*, *ABCA13*; *B*, *ABCC1*; *C*, *ABCC2*; *D*, *ABCC4*; *E*, *ABCC9*) between healthy controls (green) and asthmatics (blue). Significant ABC transporter expression differences are indicated by asterisks (*q* < 0.05 according to Mann-Whitney U test with Benjamini Hochberg correction). Both datasets (GSE4302 and GSE67472) analyzed epithelial cells from medium (3^rd^-5^th^ generation) airways and were generated from the Affymetrix Human Genome U133 Plus 2 microarray platform (using the Mas 5.0 Normalization method). Data distributions are visualized as boxplots (see Methods for additional details)

In the datasets that stratified asthmatics based on disease severity (GSE63142 and GSE76227), trends for reduced *ABCC4* and *ABCA13* gene expression were associated with increased severity of disease (**Figure 6 – Supplement Table 6**). In contrast, *ABCC1*, *ABCC2*, and *ABCC9* failed to trend with disease severity (data not shown).

**Figure 6.**
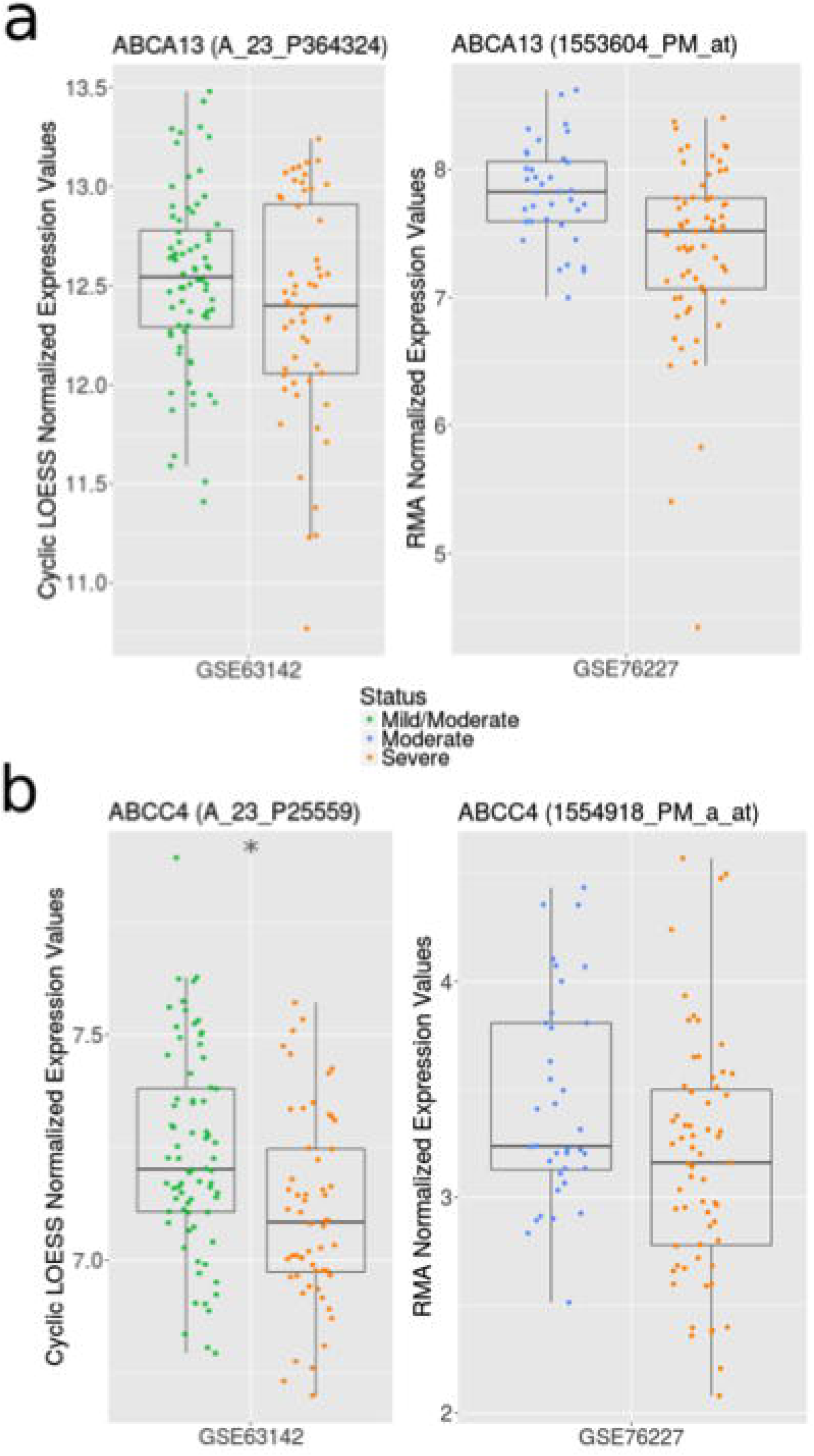
Select ABC transporter gene expression profiles associate with asthma severity. Gene expression levels of (*A*) *ABCA13* and (*B*) *ABCC4* between mild/moderate (green), moderate (blue), and severe (orange) asthmatics visualized as boxplots (see Methods for additional details). Significant ABC transporter expression differences are indicated by asterisks (*q* < 0.05 according to Mann-Whitney U test with Benjamini Hochberg correction). Both datasets used for this analysis (GSE63142 and GSE76227) analyzed epithelial cells from medium (3^rd^-5^th^ generation) airways; however, they were generated using different microarray platforms (Agilent 014850 Whole Human Genome Microarray 4×44K G4112F and Affymetrix HT HG U133 plus PM, respectively). GSE63142 uses the Cyclic LOESS Normalization method and GSE76227 uses the RMA Normalization method.

### *In vitro* validation of candidate ABC transporter gene expression changes

To validate an *in vitro* model system for interrogating candidate ABC transporters of interest, we attempted to determine if conventional cell culture systems and exposure protocols could recapitulate selected observations of differential *in situ* gene expression.

*A priori* we determined that we are unable to recapitulate a complete *in vivo* COPD and asthma phenotype *in vitro*, but are able to recapitulate the impact of cigarette smoke exposure^23^. We therefore set out to perform a cigarette smoke extract conditioned media experiment with the Calu-3 airway epithelial cell line grown under submerged monolayer conditions. We performed an exposure with cigarette smoke extract conditioned media (10% and 20%) for 24hrs and assessed *ABCA13* and *ABCC1* gene expression by quantitative-PCR. We demonstrate with our *in vitro* model that cigarette smoke extract exposure recapitulates the decrease in *ABCA13* gene expression and increase in *ABCC1* gene expression that is observed in human bronchial brushing samples (**Figure 7**) from individuals that have smoked cigarettes (**Figure 2A & 2C**).

**Figure 7.**
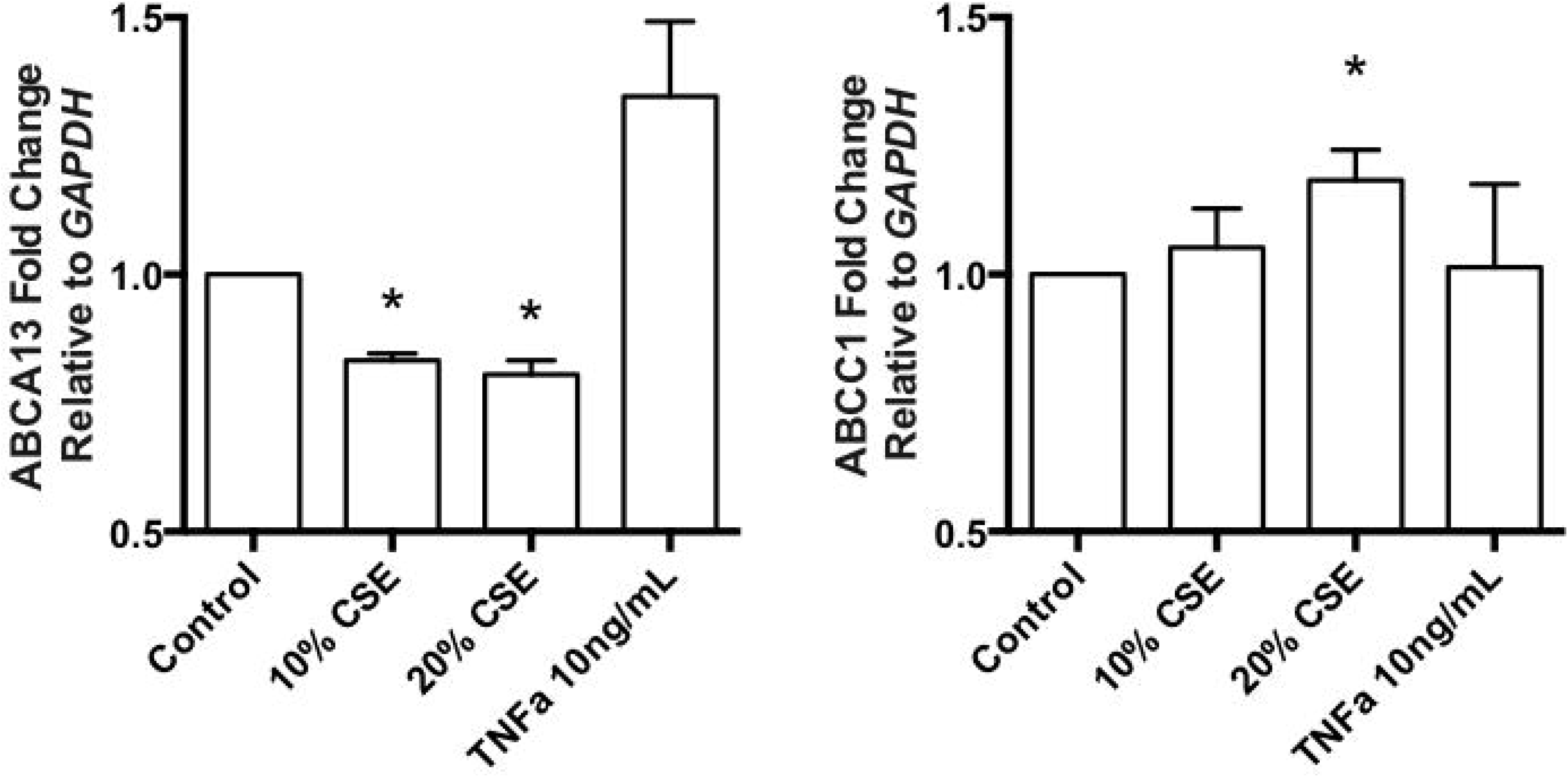
*In vitro* cell culture model systems are capable of recapitulating *in situ* observations of ABC transporter expression patterns and changes with cigarette smoke exposure. Expression levels of the *ABCA13* and *ABCC1* genes in Calu-3 cells grown under submerged monolayer conditions were assessed after exposure of cells to 10% and 20% cigarette smoke extract (CSE) conditioned media for 24 hours. Expression is relative to GAPDH levels and also compared to a negative control (unexposed Calu-3 cells) and a positive control (Calu-3 cells exposed to 10ng/mL TNF-alpha). *ABCA13* expression appears to significantly decrease as % CSE increased, whereas *ABCC1* expression increases under these conditions. n=3 * = p<0.05 relative to control.

## Discussion

The contribution of ABC transporters to respiratory mucosal immunology remains to be defined. To address this knowledge group, we performed a bioinformatic analysis of 9 distinct gene-expression datasets of primary human airway epithelial cells test our hypothesis that specific ABC transporter gene expression patterns would correlate with presence of a specific chronic respiratory disease and disease severity. The *in situ* human gene expression data demonstrate that ABC transporters are i) variably expressed in epithelial cells from different airway generations (top three expression levels - *ABCA5*, *ABCA13*, and *ABCC5*, ii) regulated by cigarette smoke exposure (*ABCA13*, *ABCB6*, *ABCC1*, and *ABCC3*), and iii) differentially expressed in individuals with COPD and asthma (*ABCA13*, *ABCC1*, *ABCC2*, *ABCC9*). Lastly, we demonstrate that an *in vitro* cell culture system of cigarette smoke exposure is amenable to investigate the consequences of differential gene expression patterns of candidate ABC transporters (*ABCA13* and *ABCC1*). Our work sets a foundation for further research into the basic biology of ABC transporters in the respiratory mucosa and suggests a potential contribution to chronic respiratory diseases.

Our analysis on ABC transporter expression profiles across airway generations was performed on samples isolated from healthy subjects at the level of the trachea, large airways (generation 2^nd^-3^rd^), and small airways (generation 10^th^-12^th^). Our data demonstrates that the expression levels of select ABC transporters are heterogeneous throughout the airway tree (e.g. *ABCC1*, *ABCC4*, & *ABCB3*) while others are homogenous (e.g. *ABCA13*, *ABCC2*, & *ABCC3*). The relative paucity of literature mechanistically describing the function of each ABC transporter in human airway epithelial cells precludes us from attributing any differential expression pattern to airway generation specific functional responses of the epithelium. Regardless, our data suggests that ABC transporter expression is differentially regulated along the airway tree, which may be important for chronic airway diseases that lead to pathology of distinct airway regions (e.g. COPD as a small airways disease^25^).

Environmental exposures including cigarette smoke, air pollution, allergens, viruses, and bacteria can induce immune mediator release in the airway epithelium^1,2^. ABC transporters may be responsible for immune mediator release, including uric acid, leukotrienes, prostaglandins, and glutathione conjugates^8,9^, that may be important for responding to environmental stimuli. To investigate possible ABC transporter-environment relationships, we explored the impact of cigarette smoke exposure and smoking cessation on *in situ* gene expression in epithelial cells. Our analyses of the impact of smoking used three datasets generated on the same microarray platform (GSE11906, GSE11784, and GSE4498)^15-17^, while our smoking cessation analysis used two additional datasets generated on two different microarray platforms (GSE37147 and GSE994)^14,18^. Conserved elevations in *ABCB6*, *ABCC1*, and *ABCC3* gene expression in response to cigarette smoke exposure were observed, accompanied by trends for decreased gene expression with smoking cessation. ABCB6 is a broad-spectrum porphyrin transporter expressed in mitochondria and plasma membranes involved in heme biosynthesis and protective responses to reactive oxygen species^26,27^. ABCB6 over-expression increases catalase gene expression and stability^28^ and is able to inhibit arsenic cytotoxicity^29^, a carcinogenic component found in cigarette smoke^30^. Despite the potential protective consequences of elevated ABCB6 expression that we observe in cigarette smokers, it is important to note that negative consequences may also arise as elevated ABCB6 levels may lead to resistance to chemotherapeutic agents^31^. Therefore, the observed up-regulation in ABCB6 is likely an important initial protective response to cigarette smoke exposure that may have untoward consequences of reducing responsiveness to chemotherapeutic agents for management of lung cancers. ABCC1 and ABCC3 also provide protective effects by transporting glutathione-conjugated anions from the cytosol to the extracellular compartment to prevent cellular accumulation of toxic metabolites^32,33^. Similar to ABCB6, ABCC1, and ABCC3 are also capable of effluxing chemotherapeutics of broad function including anti-cancer functions. Collectively it is conceivable that *ABCB6*, *ABCC1*, and *ABCC3* gene expression levels are regulated by a feedback mechanism linked to oxidative stress and toxin exposure induced by cigarette smoke exposure to help facilitate anti-oxidant activities within airway epithelial cells.

Importantly, we observe that as cigarette smokers progress to develop COPD, *ABCC1* gene expression is elevated compared to individuals that do not have COPD. Linked to cigarette smoking, elevated *ABCC1* gene expression has been observed in both non-small cell carcinoma and small-cell carcinoma lung cancers^34^. The contribution of ABCC1 biology in response to cigarette smoke exposure and to the associated development of COPD and lung carcinoma is intriguing. Our *in vitro* model of cigarette smoke extract conditioned media induction of *ABCC1* gene expression may be valuable in further interrogating this biology and the functional consequences in the context of COPD and lung carcinomas.

In contrast to elevated gene expression levels in response to cigarette smoke exposure and COPD status, we observed a conserved decrease in the expression of *ABCA13*. We also observed a decrease in *ABCA13* gene expression in asthmatics relative to healthy controls, suggesting this may be a non-specific response to an inflammatory lung environment. ABCA13 is an intriguing candidate as the expression levels are very high relative to other ABC family members, and there are no reports of endogenous or exogenous substrates, expression patterns in epithelium, or associations with respiratory mucosal immunology. Reports show associations between ABCA13 expression and cancers^35-37^, while ABCA13 variants are associated with mental health abnormalities^38,39^, although no mechanisms have been defined for either of these observations. Other ABCA family members are involved in lipid transport and dysregulation of their pathways can result in lung inflammation^40^. To gain insight into ABCA13 biology, we can examine other ABCA family members: *Abca1*-KO mice have impaired lipid transport resulting in reduced serum cholesterol and HDL, elevated intracellular lipid contents, and abnormal lung structure^41^. In contrast, mice over-expressing human *ABCA1* gene showed reduced inflammation and pathology in a model of allergic lung inflammation^42^. In humans and mice, dysfunction of ABCA3 results in surfactant deficiencies and fatal respiratory failure^12,43,44^. Based on function of these two ABCA family members in the lung, we hypothesize that ABCA13 is important in lipid handling in airway epithelial cells, and when decreased in expression, this could result in inflammation, altered surfactant production, and impaired innate immune functions. The relatively high expression levels of *ABCA13* gene throughout the airway generations and the known function of ABCA family members in lung biology warrants a further exploration of this candidate.

Our observation that cigarette smoke exposure, and to a small extent COPD status, was associated with changes in ABC transporter gene expression warranted further exploration to determine if these observations were non-specific responses to inflammatory airway environments. Although changes in ABCA13 gene expression associate with both cigarette smoking and asthma status, unique changes in ABC transporter expression profiles are observed in samples isolated from asthmatics. Elevated ABCC2 expression in asthmatics relative to healthy subjects was observed while reductions in ABCC9 were observed. ABCC2 is similar in function to ABCC1 with the shared ability to transport glutathione-conjugated xenobiotics and control intracellular oxidative stress ^45-47^. Elevations in ABCC2 that are observed in airway epithelial cells in asthmatics may aid in control of oxidative environments in the asthmatic lung and epithelial cells ^48-50^, but could also lead to extracellular transport of pharmacological agents used to control asthma due to the broad specificity of this transporter. Similar to ABCC1 in smokers, changes in ABCC2 in asthmatics may be both beneficial (management of oxidative stress) and detrimental (reduction in intracellular bioavailability of pharmacological agents). In contrast to elevations in ABCC2, we observed reductions in ABCC9 in airway epithelial cells from asthmatics. ABCC9 is also known as the sulfonylurea receptor 2 (SUR2) protein and functions as an ATP sensitive K+ channel that helps coordinate calcium levels in skeletal and cardiac muscle^51,52^. Reduced ABCC9 levels could conceivably dysregulate K+ and calcium concentrations within airway epithelial cells, which in turn could impact K+ regulated mechanisms of epithelial cell migration, proliferation, and tissue repair^53^. Irrespective of the proposed functions of these ABC transporters that are differentially expressed in epithelial cells from asthmatics, we can conclude that the different patterns observed in cells from asthmatics, smokers, or individuals with COPD suggests that regulation and function of ABC transporters is specific to a given chronic inflammatory lung disease or environmental insult.

Our bioinformatic study has some limitations that need to be recognized due to the use of publically available datasets in the NCBI GEO. The deposited datasets were developed from different microarray platforms, probesets, and epithelial cells isolated from different airway generations. Where the information was available, we have attempted to clearly disclose this in our methods and correspondingly the results sections have axes of figures labeled based on the pre-processing analysis performed by the original contributors of the datasets. Irrespective of these limitations and our inability to control for differences in microarray platform and source of epithelial cells, we emphasize that we observed common trends across independent datasets, suggesting validity of our approach and the available datasets.

In closing, we have initiated a characterization of the 48 known ABC transporters in human airway epithelial cells in the context of chronic airway diseases. Using 9 distinct gene-expression datasets of primary human airway epithelial cells, we tested our hypothesis that specific ABC transporter gene expression patterns would correlate with presence of a specific chronic respiratory disease and disease severity. It is clear that ABC transporters are i) variably expressed in epithelial cells from different airway generations, ii) regulated by cigarette smoke exposure, and iii) differentially expressed in individuals with COPD and asthma, We further demonstrate that an *in vitro* cell culture system is amenable to investigate the consequences of differential expression patterns of candidate ABC transporters, creating a foundation for further research into the basic biology of ABC transporters in lung health and disease.

## Acknowledgements

We would like to thank the individuals and research groups that have deposited their datasets to the NCBI GEO. Specific mentions of gratitude for Drs. Ian Adcock, Scott Kuo, Sally Wenzel, Prescott Woodruff, Srilaxmi Nerella for their time in addressing specific questions regarding datasets deposited in NCBI GEO. Lastly, we would like to thank Dr. Fiona Whelan for helping bring together the authors of the present manuscript for this collaboration.

